# AutophagyNet: High-resolution data source for the analysis of autophagy and its regulation

**DOI:** 10.1101/2023.03.30.534858

**Authors:** Luca Csabai, Balázs Bohár, Dénes Türei, Sowmya Prabhu, László Földvári-Nagy, Matthew Madgwick, Dávid Fazekas, Dezső Módos, Márton Ölbei, Themis Halka, Martina Poletti, Polina Kornilova, Tamás Kadlecsik, Amanda Demeter, Máté Szalay-Bekő, Orsolya Kapuy, Katalin Lenti, Tibor Vellai, Lejla Gul, Tamás Korcsmáros

**Affiliations:** Earlham Institute, Norwich, NR4 7UZ, UK; Department of Genetics, ELTE Eötvös Loránd University, Budapest, H-1117, Hungary; Heidelberg University, Faculty of Medicine, and Heidelberg University Hospital, Institute for Computational Biomedicine, Bioquant, Heidelberg, Germany; Department of Morphology and Physiology, Faculty of Health Sciences, Semmelweis University, Budapest, H-1088, Hungary; Department of Metabolism, Digestion and Reproduction, Imperial College London, London, UK; Quadram Institute, Norwich Research Park, Norwich, NR4 7UQ, UK; Department of Molecular Biology, Semmelweis University, Budapest, H-1094, Hungary; ELKH/MTA-ELTE Genetics Research Group, Budapest, H-1117, Hungary

**Keywords:** Autophagy regulation, network resource, multi-omics, signaling, big data

## Abstract

Autophagy is a highly-conserved catabolic process eliminating dysfunctional cellular components and invading pathogens. Autophagy malfunction contributes to disorders such as cancer, neurodegenerative and inflammatory diseases. Understanding autophagy regulation in health and disease has been the focus of the last decades. We previously provided an integrated database for autophagy research, the Autophagy Regulatory Network (ARN). For the last seven years, this resource has been used by thousands of users. Here, we present a new and upgraded resource, AutophagyNet. It builds on the previous database but contains major improvements to address user feedback and novel needs due to the advancement in omics data availability. AutophagyNet contains updated interaction curation and integration of over 280,000 experimentally verified interactions between core autophagy proteins and their protein, transcriptional and post-transcriptional regulators as well as their potential upstream pathway connections. AutophagyNet provides annotations for each core protein about their role: 1) in different types of autophagy (mitophagy, xenophagy, etc.); 2) in distinct stages of autophagy (initiation, elongation, termination, etc); 3) with subcellular and tissue-specific localization. These annotations can be used to filter the dataset, providing customizable download options tailored to the user’s needs. The resource is available in various file formats (e.g., CSV, BioPAX and PSI-MI), and data can be analyzed and visualized directly in Cytoscape. The multi-layered regulation of autophagy can be analyzed by combining AutophagyNet with tissue- or cell type-specific using (multi-)omics datasets (e.g. transcriptomic or proteomic data). The resource is publicly accessible at http://autophagynet.org.

## Introduction

Since its discovery nearly 60 years ago, autophagy has been widely studied to understand its mechanism, regulation, molecular function and its role in diseases. Autophagy is responsible for maintaining cellular homeostasis, differentiation, stress-response through eliminating damaged cellular organelles, misfolded proteins or intracellular pathogens. Hence, disruption in the tightly regulated molecular process leads to dysfunctional cellular processes contributing to the emergence of diseases [1]. Autophagy is induced by several internal (e.g., starvation, hypoxia or oxidative stress) and external (e.g., pathogens) factors. Most studies have focused on the regulation of the autophagic machinery through post-translational modifications (PTM), while the transcriptional and post-transcriptional regulation of autophagy are not as much uncovered [2]. To get a systems-level view and understand the concept of autophagy regulation in depth, all of these regulatory interactions should be considered and combined.

Proteins encoded by the core autophagy-related genes (ATG) operate the molecular mechanism of autophagy [3]. The organization of these proteins into complexes drives the steps of the autophagic process, such as initiation, elongation and fusion. However, there is no universal classification of these steps - some definitions are broader than others, and some proteins have overlapping functions. Discovering the regulation of the complexes involved in the individual steps can also give an insight into the workings of the autophagic machinery. Most of the literature supported data describes the process of non-selective macroautophagy, the most common form of cellular degradation. However, selective autophagic processes, such as xenophagy and mitophagy, also play important roles in maintaining normal cell function [4]. Hence, it is important to analyze both common and unique aspects of the regulation of these processes.

Regulation of autophagy through PTMs includes the phosphorylation, ubiquitination, and acetylation of autophagy proteins. Phosphorylation is responsible for protein (de)activation, such as in the case of the activation of ULK1 - a protein playing a key role in the initiation step of autophagy [5]. In contrast, the phosphorylated form of the autophagosome component LC3-II can be linked to a reduced function of double membrane formation [6]. Ubiquitination selects proteins for degradation. There is an evolutionary cross-talk between autophagy and the ubiquitin system, the core autophagy proteins mimic two ubiquitin-like protein conjugation systems (Atg12- and Atg8-conjugation systems) therefore maintaining a precise control of the process [7]. During acetylation an acetyl group is transferred to the protein causing altered features, like increased hydrophobicity or activity. In terms of autophagy regulation, under normal conditions the mTORC1 complex is acetylated, therefore the active complex can inhibit autophagy [8, 9]. Currently available large-scale post-translational databases, such as PhosphoSite [10] and PTMCode [11], do not provide filtering of data based on molecular processes, hence retrieving autophagy-related PTM information can be challenging and time consuming. Some autophagy-specific resources contain information on PTM regulation, such as THANATOS and ATdb [12, 13]. However, retrieving information from these databases to analyze omics data prove to be quite challenging as no straightforward download options are available.

Transcription factors enhance or reduce the expression of genes, including that of core autophagy proteins and their PTM regulators. With the discovery of the transcription factor TFEB, research investigating the transcriptional control of autophagy has gained increasing importance in the last decade [14], [15]. Besides, members of the Forkhead box O (FOXO) family of transcription factors have been known to play an important role in autophagy regulation in the initial phase [16]. However, current data on transcriptional regulation of autophagy genes is scattered across many resources (e.g., DoRothEA [17] and GTRD [18]), which makes it challenging to find and integrate high-quality and relevant data. To bridge this gap, TFLink [19] was developed, collecting and integrating transcription factor - target gene interactions from major databases. However, even TFLink does not provide a way to investigate direct and indirect autophagy-regulating transcription factors.

Non-coding RNAs, such as microRNAs (miRNAs) and long non-coding RNAs (lncRNAs), are key regulators of cellular processes, including autophagy. Currently, there are around 300 miRNAs described as potential autophagy regulators [20]. LncRNAs express their effect mostly through miRNA inhibition, working as miRNA sponges. Both miRNAs and lncRNAs are capable of regulating autophagy at different stages, such as at the initiation and phagophore formation [21]. With miRNA and lncRNA research advancing, there is more and more data available, though most databases still contain only predicted interactions (e.g., TargetScan [22]). As technology advances, an increasing amount of resources containing experimentally verified post-translational interactions become available. Still, they lack information on autophagy-specific aspects of these regulatory connections. There is sparse data of miRNA and lncRNA regulation of autophagy in currently available autophagy-focused resources (e.g., ATdb [13]), however most autophagy specific resources do not contain miRNA and lncRNA regulatory information at all (Supplementary Table 1).

To support systems-level analysis, the number of molecular interaction databases has been steeply increasing in the last decades, and most large interaction databases (such as Reactome [23], OmniPath [24], etc.) contain interactions of autophagy proteins and their regulators. However, in most cases these interactions are not annotated to autophagy. Besides, extracting autophagy-specific interactions from large files is challenging because these databases do not use the same structure and identifier systems, which makes it difficult to process data coming from these sources. There are a few autophagy-specific interaction databases currently available (such as Human Autophagy Database (http://autophagy.lu), Human Autophagy Modulator Database [25] and Autophagy Database [26]), but most of these resources focus only on the interactions between core autophagy proteins, or on direct autophagy PTM regulators, and many lack the information on selective autophagy.

Prompted by the lack of a detailed autophagy regulatory resource, we previously developed the Autophagy Regulatory Network (ARN) resource [27]. By combining the multiple regulation types into one multi-layered structure, the entire complex system of autophagy regulation could be analyzed. The ARN provided a robust tool to represent, visualize and analyze the complex network of autophagy regulation, and had over 7800 users from more than 60 countries in the past five years. Since its development, many advancements have been made in the field of autophagy research, with more datasets being available, including single-cell and spatial transcriptomics data. We have also received numerous user feedback and identified new, autophagy-centric datasets that should be integrated to create a more large-scale and up-to-date database tailored to the needs of the autophagy research community. Due to the significant change in the content and data types, we have created a novel resource called AutophagyNet (https://autophagynet.org). AutophagyNet integrates all the features of the ARN database as well as offers a new, extensively annotated and easy-to-use webresource containing experimentally verified protein-protein, miRNA-mRNA and lncRNA-miRNA interactions to investigate the process at a multi-layered regulatory level.

## Results

### Dataset

AutophagyNet is an integrated network resource with a multi-layered approach (Figure 1). The core of the resource is represented by 34 core autophagy proteins and their interactions. Each layer contains a type of regulator of the core machinery: i.) direct regulators have directed interactions with autophagy proteins (collected by manual curation of the autophagy literature and from autophagy-specific databases as well as large-scale protein-protein interaction databases), ii.) additional protein regulators are suggested to have interactions with direct regulators and autophagy core proteins using undirected interaction data from third party resources, iii.) transcription factors regulate the expression of members in the previous two layers and the autophagy core through directed regulatory interactions, iv.) similarly to the previous layer, post-transcriptional regulators (miRNAs and lncRNAs), are connected to the regulators in layer i-ii and the core *via* directed interactions. We have also integrated a signaling network-specific resource, SignaLink3 [28], previously developed by our group, to connect autophagy to different upstream signaling pathways. All interactions in the resource are experimentally verified and contain their primary literature source.

**Figure 1:**
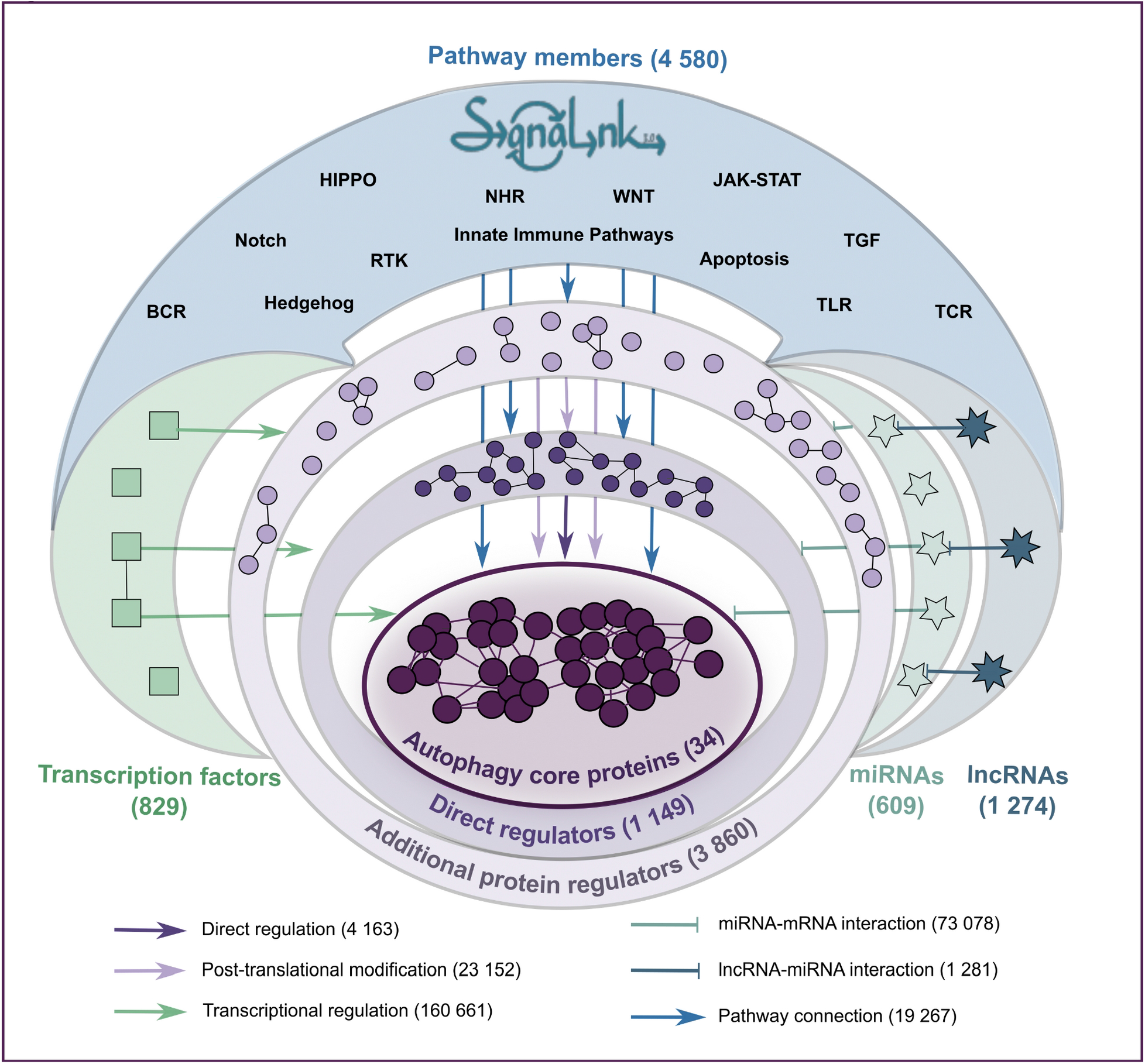
The layered structure of the AutophagyNet database. The database includes the core and regulatory proteins, transcription factors and RNAs of the process, and the diverse sets of interactions between them. Abbreviations indicate BCR - B cell receptor pathway, TCR - T cell receptor pathway, RTK - Receptor tyrosine kinase pathway, NHR - nuclear hormone receptor pathway, TLR - Toll-like receptor pathway

AutophagyNet contains over 600,000 interactions including over 19,000 unique interactors. Compared to the ARN, the number of experimentally verified interactions was increased in each layer. The number of PTM connections is also lower, as AutophagyNet only contains experimentally verified interactions, whereas ARN also included predicted interactions (Figure 2). To extend the number of interactions between the core proteins beyond manually curated interactions, we integrated interaction data between the core autophagy proteins from autophagy-specific resources (the Human Autophagy Database [25] and Autophagy Database [26]). The dataset behind AutophagyNet was made up-to-date by downloading the most recent datasets of sources used previously in the ARN resource. We have also added four further, large resources (e.g., OmniPath [24] and TFLink [19]), resulting in a total of 23 high quality resources (Supplementary Table 2). Compared to the ARN, in AutophagyNet, we have nearly doubled the number of transcription factors (from 413 to 829) and miRNAs (from 386 to 609) involved in the regulation of autophagy. With the presence of an additional layer, AutophagyNet now contains information on the effect of over 1,000 lncRNAs on autophagy, in comparison with ARN which contained data about miRNAs only. Information on the type of interaction (i.e. phosphorylation, co-localization), interaction detection methods and other annotations were also imported from the integrated resources (e.g., InnateDB [29], TFlink [19] and TarBase [30]) where available.

**Figure 2:**
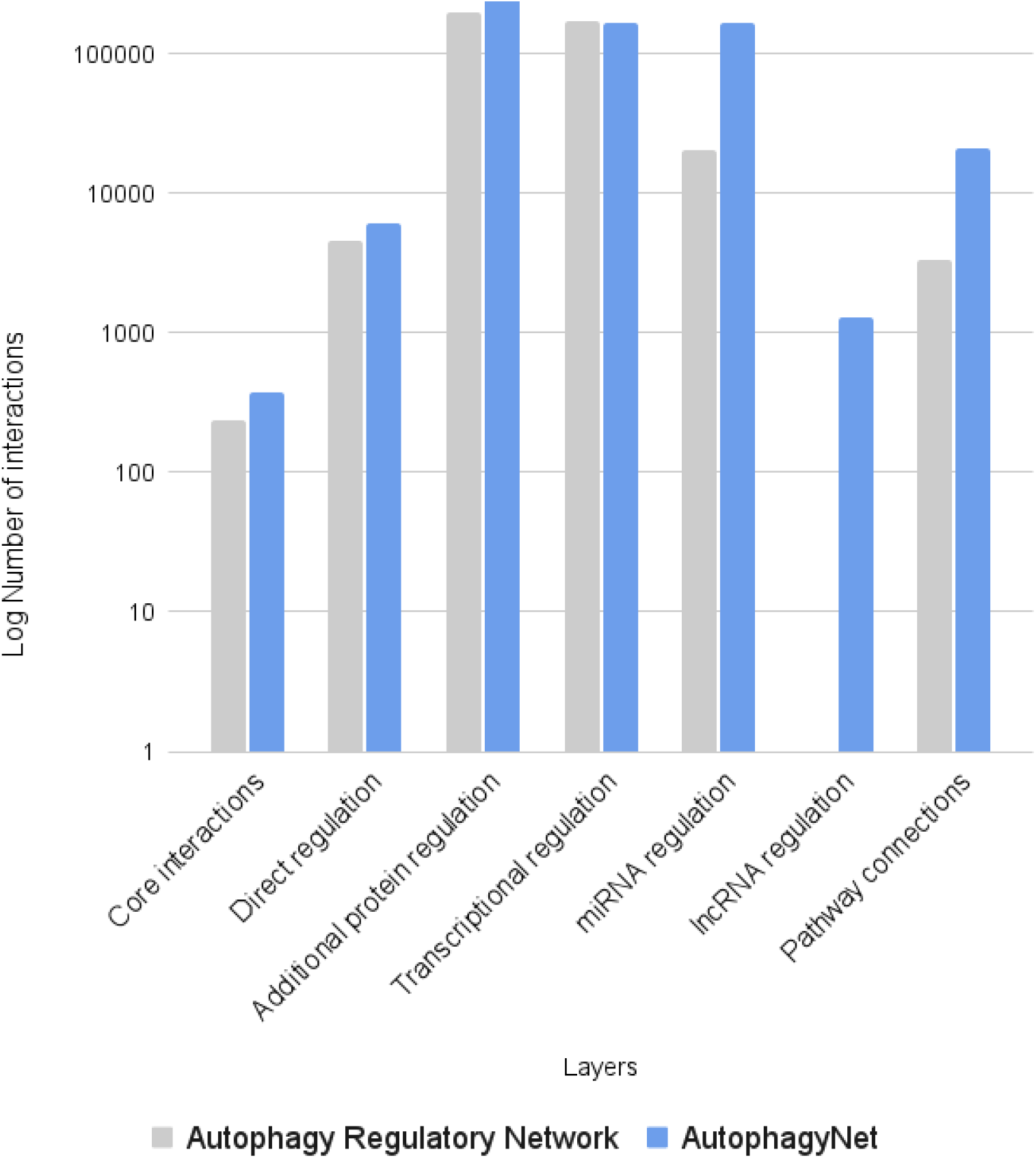
Statistics of AutophagyNet. Number of interactions are counted by layer in the previous Autophagy Regulatory Network (gray) and the new AutophagyNet (blue) resources.

### Annotations

Additional information on the properties of proteins or interactions have become key in analyzing large-scale omics datasets, as well as interpreting complex processes. Accordingly, we have included multiple general and autophagy-specific annotations to AutophagyNet. Gene/protein annotations can represent any attribute of a molecule, such as what step of autophagy they are involved in or their molecular type (e.g., enzyme, transcription factor or RNA). Further annotations can be assigned to proteins to reflect their biological role, such as which tissue they are expressed in. Interaction annotations are the features of the connection between molecules, such as directionality or the effect of the connections (i.e., activation or inhibition). The latter information is essential to develop dynamic computational models and for the modeling of drug responses for example.

As a unique feature among the existing autophagy databases, AutophagyNet contains annotations about core proteins and their direct regulators with three autophagy-specific information: i.) the phase of autophagy (e.g., initiation or autophagosome formation) the core protein is involved in (based on the Gene Ontology database [31]), ii.) the type of autophagy (e.g., macroautophagy, chaperone-mediated autophagy or xenophagy) the protein is involved in (based on Parzych KR, Klionsky DJ. (2014)[32]), iii.) the effect of regulators on autophagy activation (i.e., positive or negative) based on our manual curation. This enables the user to carry out a more focused analysis of only one step or one type of autophagy. The phase of autophagy annotation is available for all the 34 core autophagy proteins, 176 proteins are annotated with involvement of at least one type of autophagy and 710 proteins have autophagy activator or inhibitor phenotype information among core and direct regulator proteins.

Directionality and signage (activation or inhibition) of an interaction is an important aspect when analyzing regulatory interactions. These annotations are indicated in the case of every interaction, where reliable information is available from the source database. 91% of integrated interactions had known directionality information, while 26% of integrated interactions had known signage information from the source datasets. In the case of transcriptional and post-transcriptional interactions, the direction of the connections is known. However, in the case of protein-protein interactions, this is not always the case. Using a modeling approach described in the Methods section, we predicted the directionality of over 21,000 protein-protein interactions, which make up 94% of the dataset with no previous directionality information. With this method, we can identify potentially important regulators of the autophagic machinery. The sign of an interaction, which represents the effect each protein has on its direct target, is also an important aspect when trying to understand regulatory mechanisms. We predicted the signage of over 38,000 protein-protein interactions, which make up 81% of the dataset with no previous signage information (see details in the Methods section). As these predictions provide promising and crucial information for modeling the regulation of autophagy, further experimental verification is needed in order to know the certainty of these results [28].

A manual curation of autophagy literature was conducted to determine the overall impact of regulators on autophagy status. Studies that utilized gene knockout or siRNA methods to examine the effects of specific proteins on autophagy flux were selected for inclusion. Priority was given to large-scale studies that analyzed a significant number of regulators. The results of the curation were supplemented with findings from smaller-scale studies. We collected three siRNA-related studies [33–35], one study that curated data from the literature [36] and four focused studies [37–40]. In total, we identified 254 potential autophagy inhibitors, 70 potential autophagy stimulators and 429 molecules from siRNA studies whose effect was not significant on the process.

AutophagyNet contains major and minor cellular localization as well as tissue expression annotation of each protein. Using the ComPPI database [41], proteins are assigned to six cellular localizations: cytosol, extracellular, membrane, mitochondrion, nucleus and secretory pathway. This annotation makes it possible for users to filter biologically unlikely PPIs and design compartment-specific autophagy models. Tissue-specific gene expression information was combined with AutophagyNet’s network data from the Bgee database [42]. Genes can be filtered based on expression in nearly 300 tissues and 12 cell types. By providing this annotation, users can filter AutophagyNet based on their tissue of interest to create tissue-specific autophagy networks, or model pathologically significant differences in autophagy regulation in specific tissues.

### Website

The new website for AutophagyNet is available at https://www.autophagynet.org. The website was designed based on user feedback on the website of the ARN to provide a straightforward and easily accessible interface for both experimental and computational researchers.

The main page of the site presents useful information on the content and possible usage of the resource as well as links related publications regarding the database. By clicking on the feedback button, users can directly send their messages to developers *via* the pop-up window, so there is no need to look for contacts when users have a question or comment regarding the resource. The auto-completing search bar on the main page can be used to search for protein names, gene names, different identifiers or accession numbers used by standard resources (e.g., UniProt or ENSEMBL). By selecting the desired search term, the user will be navigated to the page of the selected protein. Here, they can find the detailed description of the protein with its full name as well as different accession numbers, such as UniProt, ENSEMBL and HGNC identifiers. By expanding the annotation box (Figure 3/A), information on molecule type, cellular localization and tissue expression can also be browsed. Connections of the selected protein are visualized in the network box (Figure 3/B). The network is interactive, users can zoom in and out, and move around the nodes. By clicking a node, they will be navigated to the designated page of that component. Different arrow styles represent the direction and effect of the interactions. This representation provides an easy to use tool to explore the specific network of a protein of interest without having to use any third party software. Below on the page, all interactions of the selected protein can be viewed by expanding the box for each regulation type (Figure 3/C). The source databases we used to find the interaction as well as the original publication for each interaction are annotated and linked. Therefore users can make sure that these interactions are valid and can look into some details if needed. Following user feedback, we made it possible to download directly from this page the interactions of the selected protein - this way making protein-specific research more accessible as compared to previously available options in the field.

**Figure 3:**
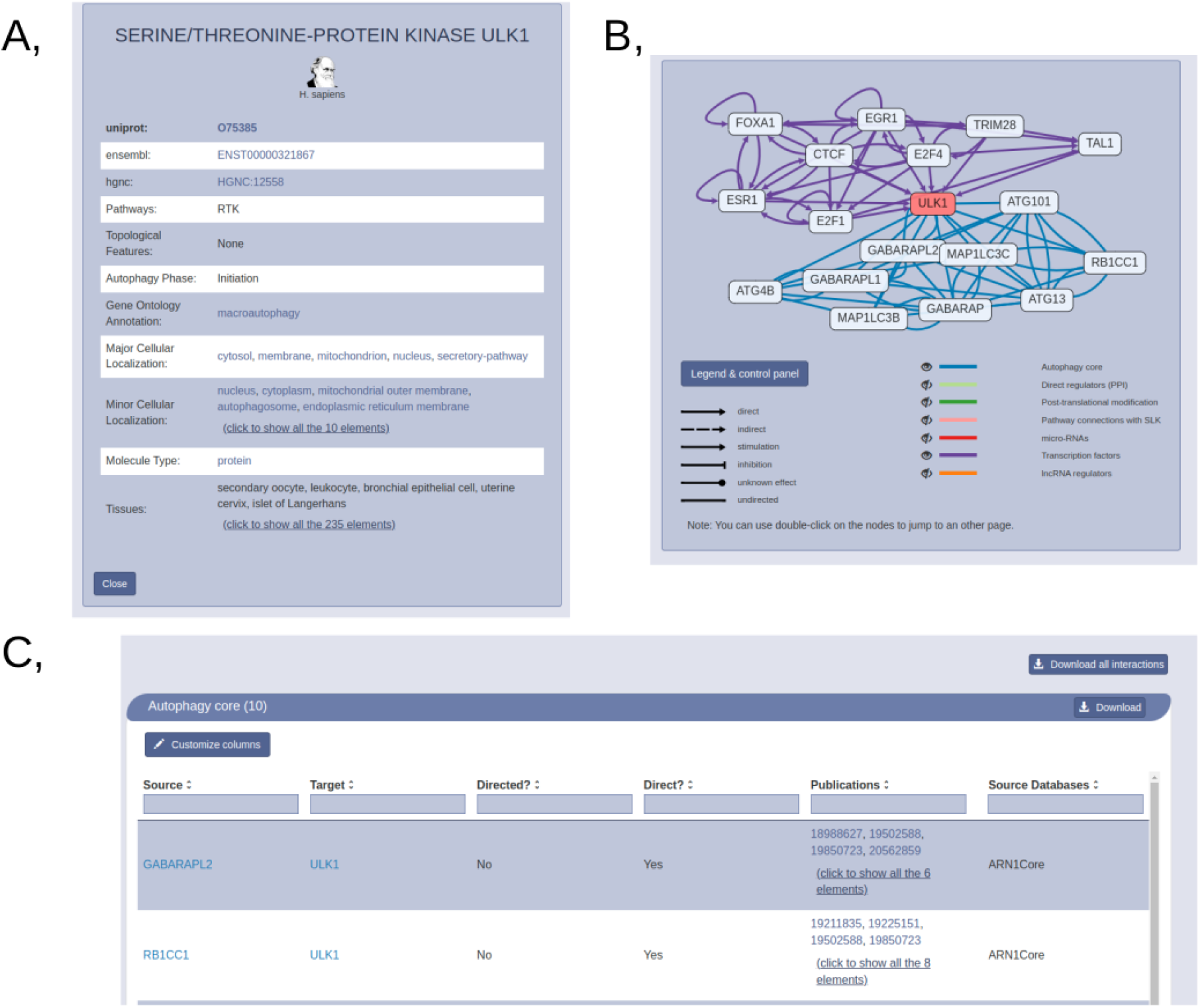
Protein page of AutophagyNet. A, Annotation box with identifiers, autophagy phase, type and tissue annotations. B, Interactive network box with core (blue) and transcriptional (purple) layers visualized. C, Download box for selected protein, annotated by source database and publications referencing the interaction.

A key feature of the AutophagyNet web-resource is that users can customize the download tailored to their specific needs (Figure 4). By navigating to the download page, users can retrieve data from the resource in just five easy steps. They can select the type(s) of regulation they are interested in and filter the dataset based on cellular or tissue localization. The core dataset can also be filtered based on the phase of autophagy and/or the type of autophagy (e.g., mitophagy) the protein is involved in. We provide a large range of widely used file formats to choose from during the download process, including simple CSV to analyze the data as text file or in Excel as well as in standardized computational file formats such as BioPAX [43] and PSI-MI Tab [44]. We also offer to download the network data directly to Cytoscape, a network visualization and analysis tool [45]. This makes it easier for non-computational researchers to visualize and analyze the retrieved data without having to work with large tables.

**Figure 4:**
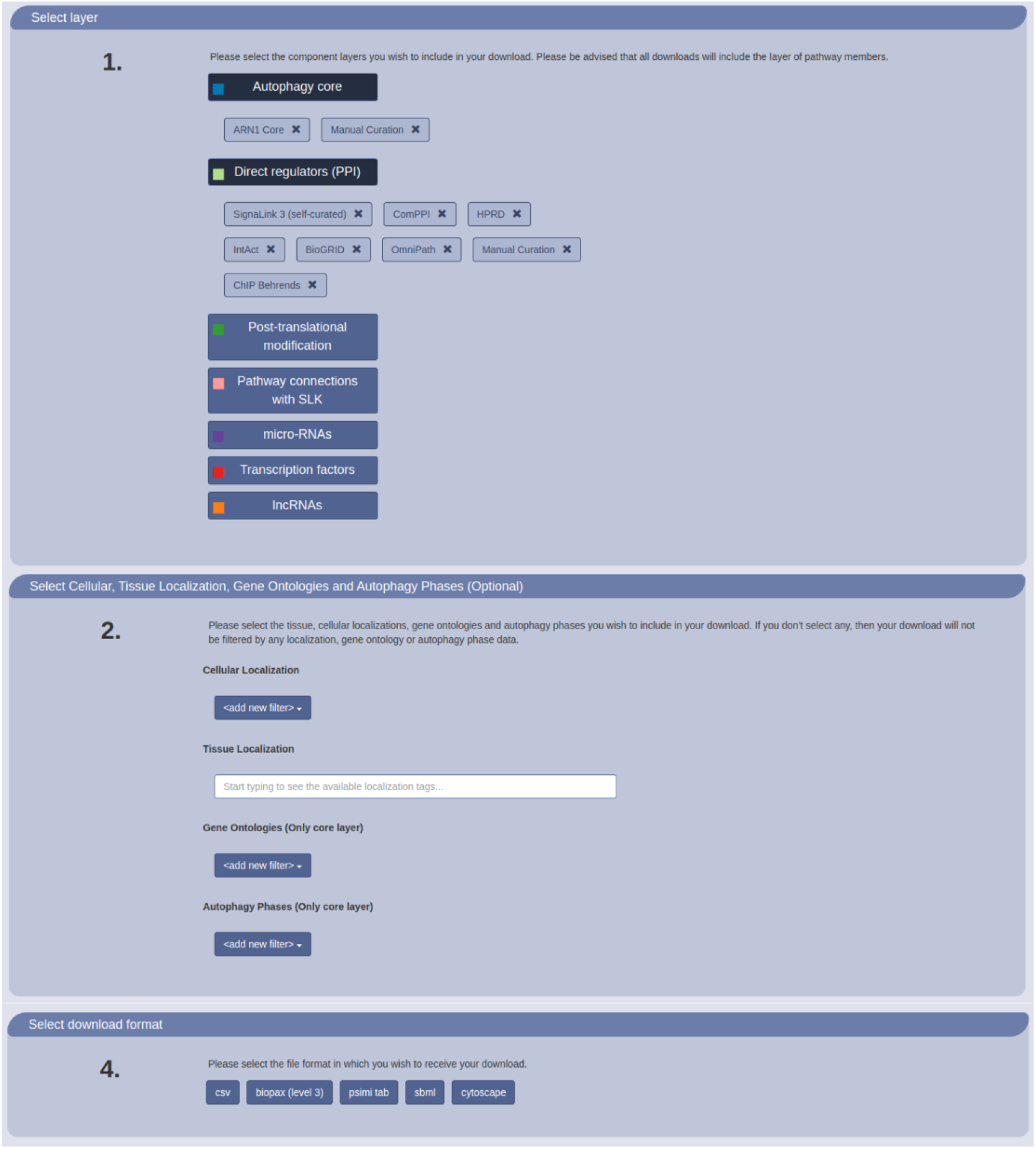
Customizable download options of AutophagyNet. Users can select regulation type, annotations and file formats to personalize the retrieved dataset on the website.

### Application

AutophagyNet can be used to analyze autophagy regulation on a systems-level and interpret autophagy-related experimental data. It provides an easy to use tool to build autophagy-specific models for prediction or data analysis purposes. A powerful aspect of the resource is that by combining it with different omics datasets, such as transcriptomics or proteomics, a multi-omics analysis of the molecular mechanisms of autophagy can be explored. Here, we present an example of how the different annotations and layers of the resource can be used for autophagy research.

AutophagyNet’s integrated annotations not only provide additional information on the function of the proteins, but make it possible for users to tailor the data of the resource for their own project. By using the customizable download filter options, AutophagyNet provides a way to explore regulation of selective autophagy, such as xenophagy, next to non-selective macroautophagy. Xenophagy is responsible for targeting invasive microbes, leading to their intracellular degradation, thereby playing an important role in antibacterial defense and immune response [46].

By selecting core members annotated to xenophagy, and selecting both direct and indirect connections of these core proteins to SignaLink3, we get an insight into which of the major signaling pathways play a role in the regulation of xenophagy. In Figure 5, we can see the xenophagy proteins, ordered based on the ‘phase of autophagy’ annotation, also retrieved from the resource. It is important to note that other core proteins can also play a role in the process of xenophagy, however based on the phase of autophagy annotations of the resource, only the presented core proteins were analyzed. We have combined regulator proteins into pathway groups based on their pathway annotation integrated from SignaLink. If one protein is involved in multiple pathways, it is counted in each group. To avoid the over enrichment of pathways containing relatively high numbers of proteins, we calculated the ratio of involved/not involved proteins in each pathway. Connections from regulator proteins to core proteins were also combined, these are represented by edges.

**Figure 5:**
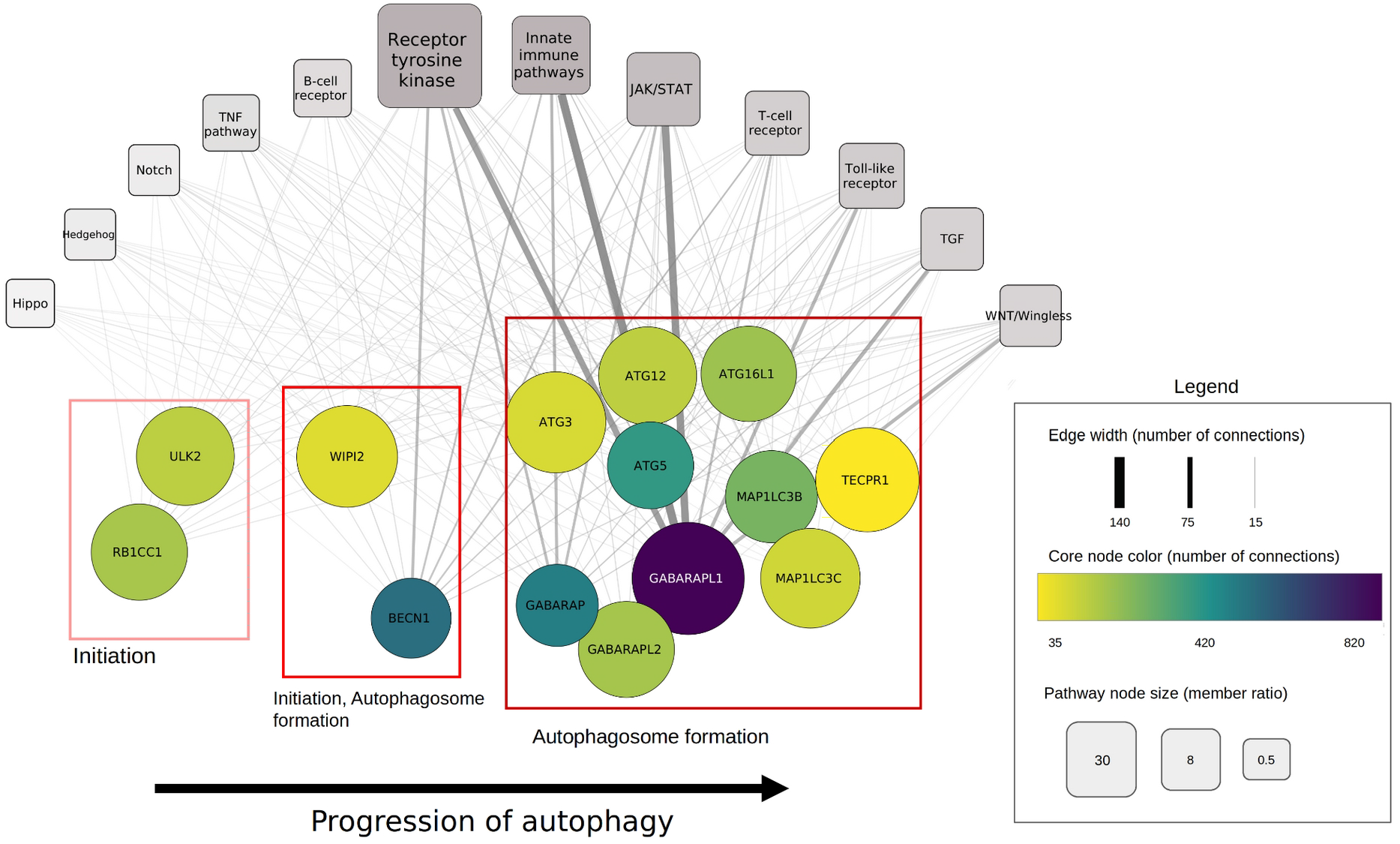
Connection of xenophagy core proteins to signaling pathways. Each circle node represents a core protein involved in xenophagy. Red boxes indicate the phase of autophagy core proteins are involved in. Rectangular nodes represent signaling pathways. The size of each pathway corresponds to the involved protein ratio, while edge width represents the number of regulatory connections from pathway members to the connected core protein.

On Figure 5, we can see that the top pathways regulating xenophagy are the Receptor tyrosine kinase (RTK), innate immune (IIP) and JAK/STAT pathways. Signaling members mostly regulate the autophagosome formation phase of xenophagy, however it is important to note that most of the xenophagy proteins are annotated to this step as well.

Based on these results, the two most regulated xenophagy members are gamma-aminobutyric acid receptor-associated protein (GABARAP) proteins and beclin 1 (BECN1). GABARAP proteins, which are major targets of many of the regulatory pathways, can interact with cell-death related proteins. Cell death acts as a suppressor of danger signals emitted by infected cells, therefore enabling an innate immune response [47]. Xenophagy represents a conserved host defense response against diverse intracellular pathogens through innate immunity [48]. The innate immune pathways are also responsible for detection and defense against intracellular pathogens, thus their connection to xenophagy can also play an important role in uncovering the mechanisms of bacterial infection.

The regulation of macroautophagy and RTK signaling are known to be closely connected as they share common vesicular trafficking pathways. RTK-autophagy cooperation has been described in multiple physiological and pathological conditions such as cancer, inflammation and aging [49]. Inhibition of the RTK pathway has also been linked to the promotion of xenophagy [50], however literature on RTK inhibitors and xenophagy is sparse. Many RTK inhibitors have been linked to a reduced bacterial load in infected organs. Inhibition of Abl-family tyrosine kinases in both *Shigella* and *Mycobacterium* infections restricts pathogen infection [51, 52]. These kinases also play a role in autophagy regulation [53], but there is no specific information of their role in xenophagy. Upon combining these findings, further analysis of the results could connect the effects of RTK inhibitors on antimicrobial defense and xenophagy. By examining the above network in a higher resolution, we can analyze exact connections between regulators and xenophagy core members. In macroautophagy, upon phosphorylation by AKT1, BECN1 bridges with 14-3-3 protein, therefore BECN1 activation is inhibited [54, 55]. Both of these connections can be seen in the network of AutrophagyNet. This indicates that this inhibition might also be present in xenophagy. A connection between the TLR pathway and xenophagy can also be seen from our example. TLR4 stimulation activates xenophagic degradation of mycobacteria *via* ubiquitination of BECN1 [56]. This connection can also be seen on Figure 5.

Using AutophagyNet, the regulation of autophagy can be examined at the post-transcriptional level, not only at the protein level. As BECN1 is an important hub of the induction of the xenophagic machinery, its post-transcriptional regulation can also reveal insights into the regulation of xenophagy. The microRNA mir-30a can suppress vesicle initiation through inhibiting *BECN1* [57]. Studies show that bacteria, such as *Mycobacterium tuberculosis* can induce expression of *mir-30a*, thereby inhibiting autophagy through *BECN1*, and promoting intracellular survival of the pathogen [58]. Further microRNA regulators of BECN1 can be found in AutophagyNet and analyzed (Figure 6), revealing similar mechanisms of pathogens downregulating xenophagy for their survival in the host.

**Figure 6:**
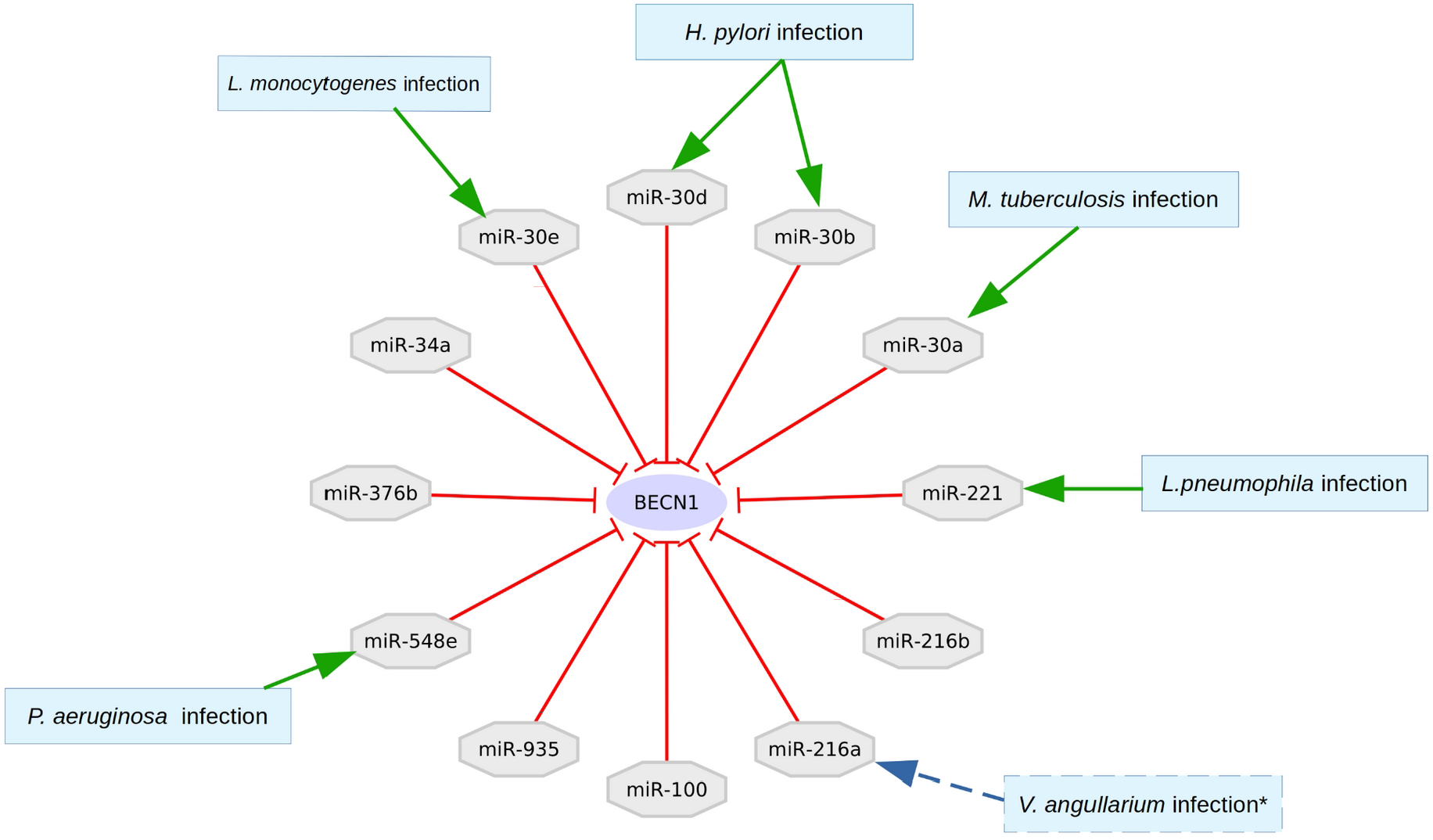
microRNA regulators of BECN1 targeted by bacterial pathogens. Bacterial infections that have been associated with microRNAs are presented. *Vibrio angillarium* infection was described in teleost fish (indicated by dashed lines), not humans. However, this could indicate the possible role of mir-216a in human bacterial infections as discussed later in the discussion section.

Other members of the mir-30 family can also be affected by bacterial infections. mir-30b and mir-30d have been found to benefit the intracellular survival of *Helicobacter pylori* [58, 59]. mir-30e is also involved in the pathogenesis of *Listeria monocytogenes*, however, this mechanism has not yet been connected with autophagy [60]. Based on our results, this pathogen could also alter the process of xenophagy to escape degradation through blocking BECN1. Figure 6 shows that mir-221 also inhibits *BECN1*. The increase of the expression of this miRNA has been linked with bacterial infection of *L. pneumophila* as well [61]. Although a direct connection between mir-221 and xenophagy has not been described, their potential connection could indicate that this microRNA plays a role in pathogen defense. mir-548e was associated with chronic *Pseudomonas* infection [62]. Low expression levels of miR-100 have also been associated with cancer, but no bacterial infection associations of the microRNA are currently known [63].

Based on the literature and these findings, analyzing the effect of interactions between xenophagy members and pathway members can provide crucial information on xenophagy regulation. This model only points out the connection between the pathways and xenophagy, but to better understand its workings, more analysis is necessary. The above application is presented to highlight various features of the resource. AutophagyNet makes it possible to analyze the regulation of autophagy and its changes under different conditions at multiple levels. Using only primary autophagy resources, this analysis would not be possible, as they only contain information on sub-sections of the complex regulatory network presented in AutophagyNet. In this case, we were able to examine how signaling pathways regulate xenophagy, but the same analysis could be carried out on transcriptional or post-transcriptional levels as well, or in a tissue specific manner. Predictions based on network models can corroborate literature findings in understanding the workings of autophagy, as well as give way to experimental validation in finding potential new drug targets in treatments of disease.

## Discussion

Here, we presented AutophagyNet, an updated and extended autophagy focused network resource. AutophagyNet is available through a user-friendly website (https://www.autophagynet.org/) improved by a customized download page tailored to researchers’ needs. AutophagyNet generates a state-of-the-art resource to study autophagy regulation. Thanks to its multi-layered nature, this resource provides the possibility to uncover different levels of autophagy regulation. The highly annotated data of AutophagyNet allows for revealing tissue-specific aspects of autophagy regulation as well as discovering the regulation of selective autophagy.

It is known that autophagy is regulated by several signaling pathways, hence the post-translational modification, especially phosphorylation, of the core machinery is a well-studied topic. While initially it was thought that autophagy is solely regulated at the protein level, a seminal report in 1999 showed that gene regulatory changes affect autophagy at a transcriptional level [64]. Moreover, autophagy regulation on the post-transcriptional level by non-coding RNAs has also been described, but less studied [65–67]. Although the knowledge on autophagy regulation has been expanded significantly, data on post-translational and (post-)transcriptional regulations are scattered across many resources - making it difficult to find and integrate high-quality and relevant data. Currently, there are large-scale databases such as BioGrid [68] TFLink [19] and TarBase [30] to describe these regulatory interactions. However, these resources do not provide autophagy regulation-related information, therefore filtering to autophagy-related data can be challenging and time consuming. To address these gaps, we developed AutophagyNet to collect the available information about autophagy regulation and organize them into multi-layered networks.

In AutophagyNet, the datasets of resources included in the ARN [27] were updated. The number of integrated resources were also extended from 19 to 32 databases (Supplementary Table 2). We also updated our manual curation by adding over 70 new connections to the core layer based on papers published between 2015 and December 2022. In AutophagyNet, we also extended the regulatory layers by including lncRNAs and their effect on miRNAs and proteins, including 1274 lncRNAs that establish 1281 interactions with miRNAs based on data from NPinter and lncRInter resources. Currently, the role of non-coding RNAs has been explored mostly in cancer, however AutophagyNet reveals several key regulatory points in autophagy through these RNA molecules. As a very needed and gap-filling feature, AutophagyNet contains multiple annotations on the autophagy proteins (role in diverse types and stages of autophagy) and on their regulators (role in the activation of autophagy). Based on feedback from the users of the ARN web-resource, we developed a new website with multiple new features (direct download from protein pages, customizable download options). AutophagyNet is a uniquely comprehensive and integrative autophagy resource that enables both computational and wet lab researchers to analyze autophagy and its multi-level regulatory layers in guiding research projects and facilitating autophagy-related biomedical studies.

There are existing, autophagy-specific databases (Supplementary Table 2) describing autophagy-related proteins, however their coverage is limited and they do not involve multi-layered regulations including both protein and RNA regulators. ADb [26] (updated in 2017) and HADb (no known date of publication or last update) describe a set of proteins involved in the process of autophagy without information on their interactions or regulatory effect on autophagy. HAMdb [25] describes interactions between autophagy proteins and their modulators including 769 proteins, 841 chemical molecules and 132 miRNAs; while AutophagySMDB [69] focuses on small molecules modulating autophagy, and describes 8 969 small molecules (information derived from the website on 26/09/2022) regulating 71 autophagy-related proteins [69]. ATdb [13] is focused on the relationship between autophagy and tumors. ATdb enlists 137 autophagy-related genes and distinguishes regulatory layers: methylation (298 enzymes), PTM (155 enzymes), miRNA (266 miRNAs) and lncRNA (658 lncRNAs) [13]. Compared to these, AutophagyNet offers a new resource to extend and unify regulatory connections of core autophagy proteins at six regulatory levels, with a total of over 600,000 connections, which is over 200,000 more than what was available in the ARN [27]. AutophagyNet also connects autophagy to signaling pathways by integrating the SignaLink3 [28] database to explore the cross-talk between them.

AutophagyNet contains a variety of annotations, providing high resolution information about regulatory interactions. Annotations describe features for molecules and their interactions. Highly annotated datasets can provide a more detailed and accurate representation of biological systems as well as cater to many different research needs. We have integrated presence/absence expression values for all proteins, which make it possible to filter AutophagyNet by tissue of interest, providing a tool for tissue-specific analysis of the regulation of autophagy. Autophagy plays a role in many diseases such as neurodegenerative diseases, inflammatory conditions and cancer [70]. Many of these pathologies are limited to one organ or tissue, hence to better understand these conditions, discovering tissue-specific workings of the autophagic machinery and its regulation is crucial. In addition, AutophagyNet includes annotations on direction and signage which help in creating a more accurate representation of regulatory connections. Annotations describing the state, type and activity of autophagy, which the core proteins and their direct regulators play a role in, facilitate the understanding of not only macroautophagy but also the less examined chaperone-mediated autophagy and different types of selective autophagy as well. By filtering the AutophagyNet resource, based on these annotations, users can customize the data for their specific research needs.

The case study shows that AutophagyNet is capable of revealing regulatory information about autophagy subtypes, such as xenophagy. By applying annotation filtering of the resource, we created a network for xenophagy regulation by signaling proteins. Based on our results we investigated literature evidence of the top three pathways regulating xenophagy. We identified immune system-related signaling pathways to regulate xenophagy, a key process involved in antimicrobial defense. Convergence of the innate immune pathway and xenophagy on cell death can highlight the importance of these regulatory connections. Although there is sparse literature-related information about the regulation of xenophagy, researches support the fact that RTK and TLR pathways interplay with autophagy as a result of sensing microbes at the cell surface [56], [50]. Connection between xenophagy and the RTK pathway has been described in the literature, however regulatory aspects of the pathway on the core xenophagy machinery still remain uncovered. By analyzing further regulatory types (e.g., inhibition by miRNAs or transcriptional regulation) available from AutophagyNet, details of the regulation of xenophagy by signaling pathways can be explored. These results could aid in understanding the pathomechanisms of diseases and identifying potential drug targets in the regulatory network.

We have demonstrated the importance of multi-level analysis of autophagy regulation through examining the post-transcriptional aspects of xenophagy regulation. The microRNA mir-30a has been widely studied in cancer, where it is an important regulator of cell proliferation and invasion, acting as a tumor suppressor [71]. Furthermore, in lung and breast cancer, the interaction between mir-30a and *BECN1* has been described as another connection between cancer and autophagy [72]. mir-30a has also been associated with drug sensitivity in gastrointestinal stromal tumors. It inhibits autophagy through *BECN1* during imatinib treatment [73]. This molecule is not only important in cancers, but in pathogen invasion as well. Bacteria such as *M. tuberculosis* can promote its survival by escaping xenophagy through inducing the expression of microRNAs [58]. Other microRNAs regulating *BECN1* have not yet been connected with autophagy in humans, however a study in teleost fish identified miR-216 as a negative regulator of bacteria-induced inflammatory responses. This mechanism works through the microRNA binding the *p62* gene, which is also a known regulator of autophagy [74]. Taken together, further experiments in human cell lines could reveal a new post-transcriptional regulator of xenophagy.

AutophagyNet also provides a tool for building autophagy-specific network models for prediction or validation purposes. Researchers can combine the network with different omics datasets, such as transcriptomics or proteomics, providing a tool to explore the molecular mechanisms of autophagy through multi-omics analysis. The multi-layered network structure of the resource offers a holistic view of biological processes.

Although autophagy is an essential function responsible for cellular homeostatic state, the current knowledge about its mechanism and regulation is poor and the information is dispersed in the literature and non-autophagy specific databases such as BioGrid or IntAct. AutophagyNet unifies datasets from manual curation, autophagy focused databases as well as other large resources, however it still does not provide a complete description of autophagy regulation. Obtaining further high-quality regulatory connections scattered across publications is challenging and time consuming, however by using text mining algorithms additional interactions could be obtained. Regulation of autophagy by small molecules is also not yet included in the resource. A future extension with metabolites and small molecule - protein interactions will allow studying how autophagy is regulated on a systems-level by microbial and host metabolites as well as by drug molecules. Supplying information on the regulatory effects of autophagy modulating drugs could aid clinical research of conditions where the dysregulation of autophagy plays a key role in the pathomechanism of disease such as IBD or Parkinson’s disease. A large amount of autophagy research is carried out in model organisms such as mice or *Drosophila melanogaster*, however available autophagy-specific regulatory interactions for these species are insubstantial. One of the potential new features is to include model organism data to make AutophagyNet available for researchers working with other widely studied species, such as mice. Integrated annotations provide crucial information for a more precise representation of autophagy regulation, however predicted annotations (such as directionality and signage) need further experimental verification to be fully trusted. As discussed above, improvements and extensions should be made to AutophagyNet as the field of autophagy research develops. The documented and automated pipeline responsible for building AutophagyNet allows us to bi-annually update the dataset, and easily add new features to the resource. This allows us to continuously better and extend the resource and cater to the specific needs of the autophagy research community.

In conclusion, AutophagyNet presented here is a powerful bioinformatics tool developed with and for the autophagy research community. On the website (https://www.autophagynet.org) post-translational, transcriptional and post-transcriptional regulation of autophagy can be examined easily and data can be downloaded in a personalized manner. Multiple annotations make it possible to create specific and accurate models of autophagy regulation. We hope this resource facilitates the process of autophagy-specific data retrieval and analysis.

## Materials and Methods

In the development of AutophagyNet, we put a big emphasis on creating an automated and well documented pipeline (https://github.com/korcsmarosgroup/AutophagyNetDB). This enables us to easily maintain the database, introduce new data in a quick and simple manner and resolve any errors efficiently.

The pipeline is made up of two main modules (Figure 7). The integration module is responsible for downloading and processing all integrated data sources (Supplementary Table 2), and converting them to our unified internal structure. This module can also be used with manually retrieved input data files as well. The integration module provides a powerful way of dealing with different input file formats and data structures, which is a common issue when dealing with datasets of different sources. Our internal structure uses SQL tables to store each individual dataset in a uniform manner, therefore in later steps of the pipeline can handle these large sets with ease. In the case of large-scale, non autophagy specific protein-protein interaction networks, we filtered the data for directed interactions, which directly targeted core autophagy proteins. These interactions, combined with processed autophagy-specific databases make up the direct regulator section of AutophagyNet. As the interactions with no known directionality might also be important regulators of the core machinery, these interactions were added to the potential regulator layer.

**Figure 7:**
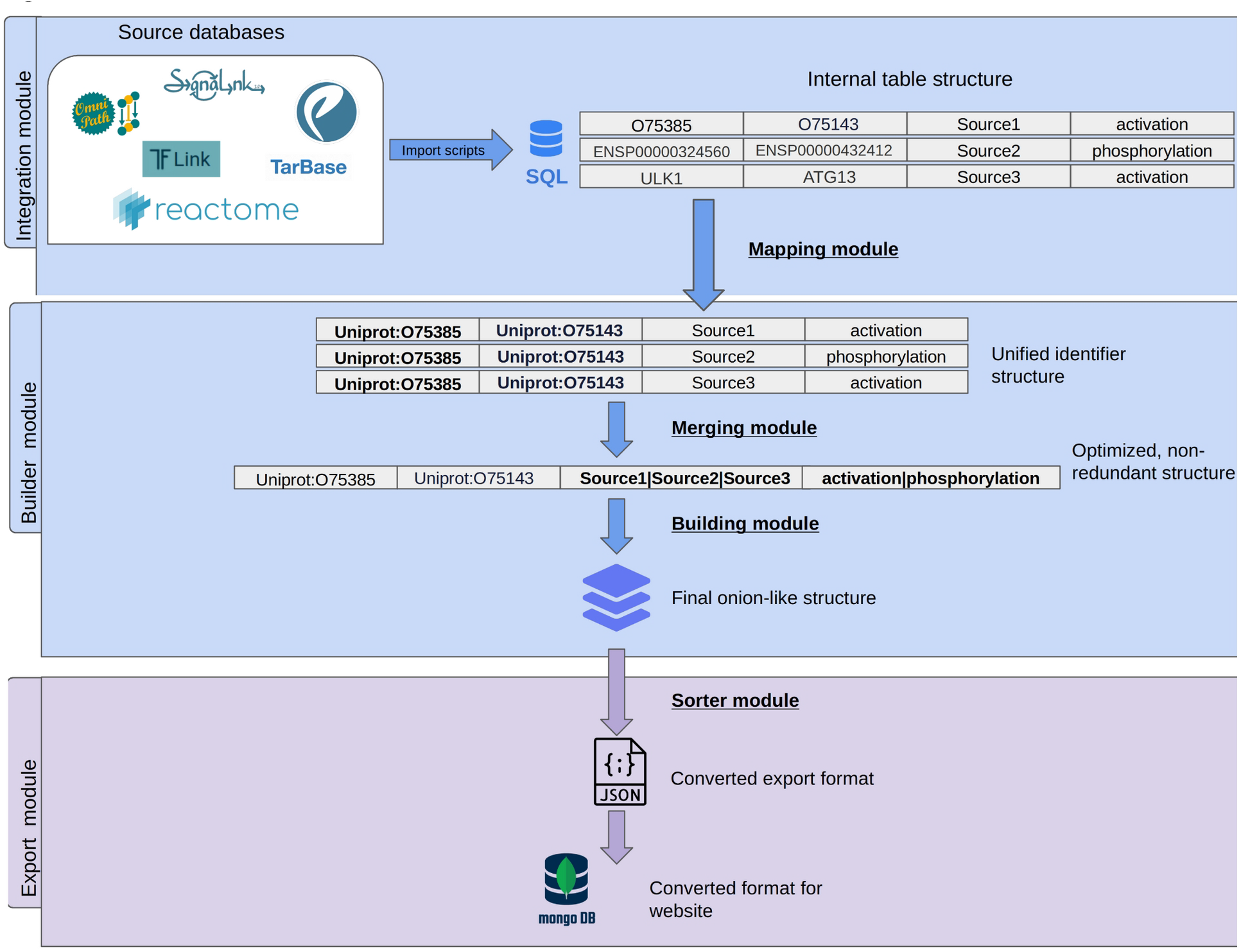
Workflow of building the AutophagyNet dataset. The pipeline is made up of two major modules: the integration and builder module. These are responsible for integrating and organizing the imported resources’ data. The export module converts the unified internal structure to file formats compatible with the user-friendly website.

The builder module consists of four smaller modules. First, each of the imported datasets are run through a mapper module, which translates the different identifiers and accession numbers to a unified format, UniProt [75] identifiers. The mapper module also assigns gene names, full protein names and cross reference identifiers from a pre-selected set of sources (i.e., IntAct [76], HPRD [77]) to each of the elements in the database. This enables us to have a unified identifier system in the resource as well as provides users with the option of searching based on a multitude of identifier types. In the merger module, each layer of the mapped data is put through, which makes sure there are no redundancies in the data. If an interaction is present multiple times in our dataset, with different annotations (i.e., coming from a different source database, having a different publication), the interactions will be merged into one entry, with all annotations combined (Figure 5). This way there is no data loss and no duplication of interactions. When each layer is mapped and merged, the onion-like multi-layered network is created by looking for overlaps in interactions between the separate layers by the building module. Afterwards, predictions (direction and effect) and annotations described in previous sections are added. Direction prediction was carried out by methods from our previous work [28], which built on approaches of Liu et al. and Rhodes et al. [78, 79]. Signage prediction methods also from our earlier work [28] were applied. This method used an RNAi screen based prediction method developed by Vinayagam et al. [80]. The calculated scores of the predictions are added to the confidence score column of each interaction in the internal SQL table. Major and minor cellular localization data was integrated from the ComPPI [41] database. Tissue expression data was integrated by using presence/absence information from the Bgee database [42]. Both the above annotations are added to the record of each node. Finally in the export module, the internal SQL structure is converted into the final json format, which is the final format required by the mongoDB database management program (https://www.mongodb.com/) running behind the website.

## Acknowledgements

We thank the comments and feedback of the users of Autophagy Regulatory Network and the pilot of the AutophagyNet web resources as well as the suggestions and ideas from the past and current members of the LINKgroup, VellaiLab, and KorcsmarosLab (previously NetBiol group). L.G., M.M. and M.P. were supported by the BBSRC Norwich Research Park Biosciences Doctoral Training Partnership (grant numbers BB/M011216/1 and BB/S50743X/1). The work of T.K. and D.M. was supported by the Earlham Institute (Norwich, UK) in partnership with the Quadram Institute (Norwich, UK) and strategically supported by a UK Research and Innovation (UKRI) Biotechnological and Biosciences Research Council (BBSRC) Core Strategic Programme Grant for Genomes to Food Security (BB/CSP1720/1) and its constituent work packages, BBS/E/T/000PR9819 and BBS/E/T/000PR9817, as well as a BBSRC ISP grant for Gut Microbes and Health (BB/R012490/1) and its constituent projects, BBS/E/F/000PR10353 and BBS/E/F/000PR10355. L.G., M.O. and T.K. are supported by the Division of Digestive Diseases at Imperial College London that receives financial and infrastructure support from the NIHR Imperial Biomedical Research Centre (BRC). T.V. was supported by the Hungarian National Research, Development and Innovation Office (K131458, PD131839 and K132439, correspondingly) and by the ELKH/MTA-ELTE Genetics Research Group (01062). The authors acknowledge funding and support from the UKRI BBSRC Core Capability Grant (BB/CCG1720/1) and the work delivered via the BBSRC National Capability in e-Infrastructure (BBS/E/T/000PR9814) at the Earlham Institute by members of the e-Infrastructure group. D.T. was supported by the German Federal Ministry of Education and Research (Bundesministerium für Bildung und Forschung BMBF) Computational Life Sciences LaMarck (grant no. 031L0181B). O.K. was supported by NKFIH FK-134267 (National Research, Development and Innovation Office, Hungary).

## Abbreviations

ARN: Autophagy Regulatory Network
ATG: autophagy-related genes
BCR: B cell receptor pathway
BECN1: beclin 1
GABARAP: gamma-aminobutyric acid receptor-associated protein
IIP: innate immune pathway
lncRNA: long non-coding RNA
miRNA: microRNA
NHR: nuclear hormone receptor pathway
PTM: post-translational modification
RTK: receptor tyrosine kinase pathway
TCR: T cell receptor pathway
TLR: Toll-like receptor pathway

