## Supplementary Table 1, Supplementary Table 2 for "AutophagyNet: High-resolution data source for the analysis of autophagy and its regulation"

**Supplementary Table 1 -** Comparison of available autophagy specific resources with AutophagyNet

| | AutophagyNet | HAdb | THANATOS | HAMDB | ATdb | Autophagy SMDB |
|---|---|---|---|---|---|---|
| **Contains manual curation** | Yes | Yes (based on literature curation) | Yes (based on literature curation) | Yes | Yes | Yes |
| **Contains integrated data** | Yes | No | No | No | Yes | Yes (FDA approval status) |
| **PPIs** | | No interactions, 232 proteins | 4237 | No interactions (796 proteins) | | Small molecules regulating 71 proteins |
| **PTMs** | Yes | No | Yes (PTM sites) | No | Yes (155) | No |
| **miRNAs** | Yes | No | No | Yes (132) | Yes (658) | No |
| **lncRNAs** | Yes | No | No | No | Yes (266) | No |
| **Interactive view** | Yes | No | No | No | Yes | Yes |
| **Customizable download** | Yes | No | No | Yes (multiple options/file formats) | Yes*limited | |
| **Search whole database** | Yes | Yes | Yes | Yes | Yes | Yes |
| **Species** | Human | Human | Human + 7 model organisms | Human | Human | Mammalian cells or in vivo |
| **Localization information** | Tissue and cellular localization | No | No | Cell filtering | Tissue and cell localization | Tissue and cell localizations |

**Supplementary Table 2** - Description of data sources used in AutophagyNet

| Resource name | Description | Reference | URL |
|---|---|---|---|
| Autophagy Regulatory Network | Previous version of current resource. | (Türei et al, 2015) | http://autophagyregulation.org/ |
| HADb | Complete and an up-to-date list of human genes and proteins involved directly or indirectly in autophagy as described in literature. | | http://autophagy.lu/ |
| Behrends et al | Proteomic analysis of the autophagy interaction network in human cells under conditions of ongoing (basal) autophagy, revealing a network of 751 interactions among 409 candidate interacting proteins with extensive connectivity among subnetworks. | (Behrends et al, 2010) | https://www.nature.com/articles/nature09204 |
| Self-curated interactions | | | |
| COMPPI | A database of cellular compartment specific protein-protein interactions | (Veres et al. 2014) | https://comppi.linkgroup.hu/protein_search |
| IntAct | Database mostly containing PPI data, annotated by IMEx standards, containing detailed description of experimental conditions of the interactions. | (Del Toro et al. 2022 | http://www.ebi.ac.uk/intact/ |
| OmniPath | Literature curated mammalian signaling pathways | (Turei et al, 2016) | http://omnipathdb.org/ |
| HPRD | Human protein reference database, containing PPIs and posttranslational modifications curated from experiments supplied with their types annotated. It also contains information about upstream enzymes responsible for protein modifications and alternative subcellular localization of the described proteins. | (Peri et al. 2003) (Keshava Prasad et al. 2009) | http://www.hprd.org/ |

| | | | |
|---|---|---|---|
| BioGrid | BioGRID is a biomedical interaction repository with data compiled through comprehensive curation efforts. Accessible in tabular format with UniProt IDs and PubMed references. | (Stark et al. 2006) (Breitkreutz et al. n.d.; Stark et al. 2011) | http://thebiogrid.org/ |
| PTMCode2 | Resource of known and predicted interactions between protein post-translational modifications and interacting proteins. | (Minguez P et al, 2012) | https://ptmcode.embl.de/ |
| PhosphoSite | Combination of low-, and high-throughput data sources of phosphorylation sites in human, mouse and other species. | (Hornbeck et al. 2004) (Hornbeck et al. 2012) | http://www.phosphosite.org/homeAction.do |
| Ramirez et al. | Potential scaffold proteins in interactomes. | (Ramirez et al. 2009) | https://www.cell.com/trends/cell-biology/fulltext/S0962-8924(09)00271-2?_returnURL=https%3A%2F%2Flinkinghub.elsevier.com%2Fretrieve%2Fpii%2FS0962892409002712%3Fshowall%3Dtrue |
| SignaLink3 | A database assigning proteins to signaling pathways using the full texts of pathway reviews. Compared to most signaling resources, SignaLink uses more than 20 review papers per pathway on average. It aims at reducing data and curation errors during the curation process. | (Csabai et al. 2022) | http://signalink.org/ |
| TFLink | Provides comprehensive and highly accurate information on transcription factor - target gene interactions, nucleotide sequences and genomic locations of transcription factor binding sites for human and six model organisms. | (Liska et al, 2022) | https://tflink.net/ |

| | | | |
|---|---|---|---|
| TarBase | Interaction database of experimentally validated miRNA-target interactions. | (Chou CH et al, 2018) | http://mirtarbase.mbc.nctu.edu.tw/php/index.php |
| miRecords | Resource for animal miRNA-target interactions. Validated targets from literature curation of experimentally validated data. | (Xiao F et al, 2009) | http://c1.accurascience.com/miRecords/ |
| mirDeathDB | An integrated database containing experimentally identified miRNAs and their targets in the PCD network. | (J Xu et al, 2012) | http://www.rna-world.org/mirdeathdb |
| mir2Disease | Manually curated database providing information of miRNA deregulation in human diseases. | (Jiang Q et al, 2009) | http://watson.compbio.iupui.edu:8080/miR2Disease |
| lncRInter | Reliable and high quality lncRNA interaction database containing experimentally validated data extracted from peer-reviewed publications. | (Chun-Jie Liu et al, 2017) | http://bioinfo.life.hust.edu.cn/lncRInter/ |
| miRSponge | Database for understanding transcriptome communication of different RNA classes (miRNA, lncRNA) | (Wang et al. 2015) | http://bio-bigdata.hrbmu.edu.cn/miRSponge/ |
| NPInter | An integrated database of noncoding RNA interactions. | (Xueyi Teng et al, 2020) | https://www.bioinfo.org/NPInter |
| StarBase | Collection of interaction networks of lncRNAs and miRNAs | ( Li et al., 2014) | http://starbase.sysu.edu.cn/starbase2/index.php |
